# PanPA: generation and alignment of panproteome graphs

**DOI:** 10.1101/2023.01.19.524778

**Authors:** Fawaz Dabbaghie, Sanjay K. Srikakulam, Tobias Marschall, Olga V. Kalinina

## Abstract

**Motivation:** Compared to eukaryotes, prokaryote genomes are more diverse through different mechanisms, including a higher mutation rate and horizontal gene transfer. Therefore, using a linear representative reference can cause a reference bias. Graph-based pangenome methods have been developed to tackle this problem. However, comparisons in DNA space is still challenging due to this high diversity. In contrast, amino acids have higher similarity due to evolutionary constraints, resulting in conserved amino acids that, however, may be encoded by several synonymous codons. Coding regions cover the majority of the genome in prokaryotes. Thus, building panproteomes leverages the high sequence similarity while not losing much of the genome in non-coding regions.

**Results:** We present `PanPA`, a method that takes a set of multiple sequence alignments (MSAs) of proteins or protein clusters, indexes them, and builds a graph for each MSA. In the querying step, it can align DNA or amino acid sequences back to these graphs. We first showcase that `PanPA` generates correct alignments on a panproteome from 1,350 *E. coli*. To demonstrate that panproteomes allow longer phylogenetic distance comparison, we compare DNA and protein alignments from 1,073 *S. enterica* assemblies against *E. coli* reference genome, pangenome, and panproteome using `BWA`, `GraphAligner`, and `PanPA` respectively, where `PanPA` was able to produce around 22% more alignments. We also aligned DNA short-reads WGS sample from *S. enterica* against the *E. coli* reference with `BWA` and the panproteome with `PanPA`, where `PanPA` was able to find alignment for 69% of the reads compared to 5% with `BWA`

**Availability:** `PanPA` is available at https://github.com/fawaz-dabbaghieh/PanPA

**Supplementary information:** Supplementary data are available at *Bioinformatics* online.

## 1 INTRODUCTION

Prokaryotes have been living on Earth for billions of years, during which they continued to evolve rapidly. With the geochemical changes on the planet, bacteria needed to adapt in order to survive these environmental and habitat changes, which led to their vast genetic diversity (Dunlap, 2001). Looking at stable environments like garden soil, lakes, or coastal seawater which do not have extreme environmental changes, we observe a large diversity in their content of prokaryotes; and it is expected that not more than 1% of the bacteria in these samples actually grow in laboratory cultures (Amann *et al*., 1995), which suggests that the true diversity is even larger. It has been estimated that the total number of prokaryotic cells on Earth is around 4 − 6 × 10^30^ and their cellular carbon amount is 3.5 − 5.5 × 10^14^ kg (Whitman *et al*., 1998).

With the fast development of sequencing technologies, and, as a consequence, the fast production of large amounts of sequences, the diversity, and variability of prokaryotic genomes have become even more apparent (Perna *et al*., 2001). One way to understand new genomes and their diversity is by comparing their DNA to some well-studied reference genomes of the same species. Therefore, sequence alignment has been a cornerstone in bioinformatics for many years: it is extremely useful for finding homology between genes and protein families, finding conserved regions, understanding evolutionary relationships between organisms through their genome sequences, and many other important functionalities (Higgins, 2001).

In many cases, sequencing reads of a new sample are directly analyzed by comparing them to a reference genome, that is, to one genome representative of the species. However, the linearity of a reference genome can lead to biases, for example, if our sequence contains a non-reference allele which leads to incorrect or missing alignments (Chen *et al*., 2021). These effects are more pronounced in highly-variable organisms like bacteria. To describe this genomic variability, the terms *core* and *accessory* genes were first coined by Tettelin *et al*. (2005), where the “core” genes refer to essential genes (e.g. housekeeping genes) that are found in all or nearly all isolates, and the “accessory” genes (sometimes called “dispensible” genes) refers to the genes that are not present in every genome or isolate sequenced. The term *pangenome* was first mentioned by (Sigaux, 2000) describing a database of tumor genome and transcriptome alterations, as well as relevant normal cells.

In recent years, graph representations of pangenomes have become more widespread, providing a more complete picture of pangenomes than a simple distinction into core and accessory genes. In graph-based models of pangenomes, one represents the genomic variability of a population in a graph data structure where nodes are labeled with sequences and edges represent sequences that are adjacent to each other in one or more genomes in a population (Eizenga *et al*., 2020). One can then use these graph data structures as a reference instead of using a linear reference to reduce biases (Paten *et al*., 2017), which entails the need to align sequences to a graph. Sequence alignment and pattern matching to a string graph is not a new problem; it has been described almost three decades ago. Pioneering studies include (Manber and Wu, 1993) where pattern matching on hypertext was described and (Akutsu, 1993) where an algorithm for exact pattern matching to a hypertext in a tree structure was developed. In 1995, Park and Kim (1995) described regular pattern matching on a directed acyclic graph (DAG). Later on, an algorithm that does pattern matching on *any* hypertext was developed by Amir *et al*. (2000), then Navarro (2000) improved both time complexity and space complexity.

In 2002, similar algorithms were applied to biological data by Lee *et al*. (2002) where they describe the use of the algorithm Partial Order Graphs in constructing MSAs and aligning to the graph representation of the MSA using dynamic programming (DP). In essence, it is a modified version of the common sequence alignment dynamic programming algorithms, where we also need to take care of the incoming edges connecting a certain letter in the graph while calculating the cell’s score. In recent years, more tools have been implemented to perform sequence-to-graph alignments with better speeds and accuracy (Rautiainen and Marschall, 2020; Sirén *et al*., 2021; Ivanov *et al*., 2020).

So far, all tools for pangenomes have mostly been implemented to be used for different samples or strains in a single species: in bacteria, for example, for *E. coli* (Colquhoun *et al*., 2021), in plants, for *Cucumis sativus* (Li *et al*., 2022), and in humans (Li *et al*., 2020; Eizenga *et al*., 2020), including the work of the Human Pangenome Reference Consortium (HPRC) (Liao *et al*., 2022). Due to the high diversity in bacteria, these tools typically cannot be used for inter-species comparisons at the DNA level, as the diversity is too high to make meaningful alignments. This problem is even more exacerbated in highly-diverse and less-studied clades like *Myxobacteria* or *Actinomycetes*, for example, which are an important source of natural products that can be used in drug discovery (Gerth *et al*., 2003). The diversity in these clades is much higher than what is already described due to limitations in cultivation and in-lab growth (Mohr, 2018).

In these cases, however, one can still trace the sequence similarity by switching to amino-acid residues alignments, i.e. looking only into coding regions, as these alignments will have a higher quality compared to DNA sequence alignments, due to several reasons. First, amino acid sequences are evolutionary more conserved compared to the total genome DNA sequence, as proteins have a specific biological function. Moreover, as the amino-acid alphabet is larger, the “signal-to-noise ratio” is better (Wernersson and Pedersen, 2003). Second, some of the errors introduced during sequencing can cause a frameshift during alignments, which can be avoided when using amino acids (Sheetlin *et al*., 2014). Third, in amino-acid sequence alignment, we usually use a substitution matrix instead of just edit distance in DNA sequence alignment, better capturing biological reality (Bininda-Emonds, 2005). In prokaryotes, the fraction of non-coding regions in the genome can range from 5 to 50%. However, for the vast majority, the fraction is less than 18% (Rogozin *et al*., 2002), further motivating a focus on coding sequences.

Here, we propose a new tool `PanPA` to conduct pangenomic analyses with respect to protein sequences. `PanPA` allows building directed acyclic graphs for each individual protein or protein cluster in order to represent a pangenome. Computing alignments in amino acid space can give a big advantage in terms of finding more sequence similarity and being able to align more phylogenetically distant organisms against each other while losing relatively little genome information. Westbrook *et al*. (2017) showcased how aligning in protein space introduces significant improvements in alignment accuracy and functional profiling in a metagenome scenario.

The idea of having many graphs representing a pangenome instead of one large graph was presented in Colquhoun *et al*. (2021); in their tool `Pandora`, the authors define a pangenome as a collection of *local graphs* where each local graph represent some locations in the genome that can be pre-defined by the user.

`PanPA` combines the two ideas of i) having a pangenome consisting of many smaller graphs, where each graph represents a protein or a protein-cluster, and ii) working in amino acid space rather than nucleotide sequences to support pangenomic analyses over larger evolutionary distances. We call such a collection of graphs a *panproteome*. We showcase the utility of `PanPA` by performing alignments of proteins and raw short reads from *Salmonella enterica* assemblies against a *E. coli* panproteome.

## 2 METHODS

The idea behind `PanPA` is that we want to build a panproteome of a collection of protein sequences or protein clusters. In this definition of a panproteome, each protein or protein cluster is represented as a separate graph. Therefore, our pipeline starts from multiple sequence alignments (MSAs) provided as input, where each MSA represent one protein or a cluster, and the pipeline goes through three major steps:

1. Building an index from the input MSAs.
2. Constructing a directed graph from each MSA.
3. Aligning query sequences to these constructed graphs with the help of the index constructed from these MSAs.

### 2.1 Building index from MSAs

For each sequence *N* of length *m*, we define a substring *s* = *N* [*i, j*], where 0 ≤ *i* ≤ *j m* ≤ 1, as a substring of *N* starting at position *i* and ending at position *j* with the length of *j − i* + 1. A *k*-mer from a string *N* is then defined as a substring of length *k*. We also define a function min(*S*) that takes the set *S* = {*s*_1_, *s*_2_, …, *s*_*w*_} of size *w* containing *w* equally-lengthed strings and returns the lexicographically smallest string in this set; we call this function a *minimizer*.

To construct a *k-mer based index*, for every string *N*, the seeds extracted from that string form a set *S*_*seeds*_ comprising every consecutive *k*-mer from *N*. *S*_*seeds*_ = {*N* [0, 0 +*k* − 1], *N* [1, 1+*k* 1], …, *N* [*i, i*+*k* − 1] }; ∀*i* {0, …, (*m* − *k*)}, where each string *N* of length *m* will contain (*m* − *k*+1) *k*-mers.

A *minimizer based index* was originally developed by (Schleimer *et al*., 2003) and was first used in bioinformatics to reduce storage requirements for sequencing data by (Roberts *et al*., 2004). In this approach, for each sequence *N*, instead of taking the set of all consecutive *k*-mers as seeds, we take the set *S*_*seeds*_ that contains the minimizer of every consecutive window of *w k*-mers, i.e. we take the smallest seed in a set of seeds for each consecutive window of seeds. *S*_*seeds*_ = {min(*S*_0,*w*_), min(*S*_1,*w*_), …, min(*S*_*i,w*_)}; ∀*i* ∈ {0, …, (*m* − *w* − *k* + 1) }, where *S*_*i,w*_ is a set of *w* consecutive *k*-mers starting at position *i* in the string *N*.

In `PanPA`, both a *k*-mer based and a minimizer-based index is implemented and can be used alternatively. In both cases, the index stores a key-value map, where the keys are the set of all *k*-mers or (*w, k*)-minimizers extracted from each sequence in the input MSAs, and the values are ordered lists of MSAs where that key was found, the ordering of the values is based on the number of times that key showed up in that certain MSA. More on the indexing detail is described in “Implementation”.

### 2.2 Generating a Directed Acyclic Graph from an MSA

For this step, we developed a simple algorithm to turn each MSA into a graph in the *graphical fragment assembly* (GFA) format, where each original sequence from the MSA is represented in the GFA as a path. This algorithm runs in 𝒪 (*n* × *m*) time where *n* is the number of sequences in the MSA and *m* is the length of the alignment and has two steps: (1) generating the graph, (2) compacting the graph.

#### 2.2.1 Generating the graph

We define the alphabet *A* as the amino acid alphabet, and the matrix *M* = (*a*_*i,j*_) ∈ {*A* ⋃ −} ^*m×n*^, each column in matrix *M* is a vector {*A* ⋃ −} ^*n*^ and each row is a vector {*A* ⋃ −}^*m*^.

In a nutshell, the algorithm loops through each column vector at position *j* where 0 ≤ *j* ≤ *m*−1, and for each of these vectors, it constructs a new node *node*_*j*_ (*c*) for each unique character *c* ∈ *A*. Edges are then added between two nodes 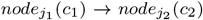 (where *j*_1_ *< j*_2_) if and only if the characters *c*_1_ and *c*_2_ were consecutive in one of the rows in matrix *M* after ignoring the character {−}.

In Figure 1 we have an MSA with 3 sequences, in the figure we have what we call a *current* column (in yellow), and *previous* column (in red), for each iteration through the columns, the algorithm goes through each character in that column, if the character is new then a new node is initialized in that step for that character (lines 18-22 in Algorithm 1), otherwise, the character is not new, then we assign that character the corresponding node identifier that was already constructed for the same character. After building nodes from a column *j*, we synchronize with *j* − 1 (if it exists) (lines 2-10 in Algorithm 1), where we go through each row *i* in both columns, and for every row *i* we have three choices: (1) if *c*_*i,j*_, *c*_*i,j*+1_ ∈ {−} (e.g. first two gaps in the second sequence in Figure 1) then there is nothing to do; (2) if *c*_*i,j*_, *c*_*i,j*+1_ ∈ *A*, then we need to draw an edge between *node*_*j*_ (*c*_*i,j*_) and *node*_*j*+1_(*c*_*i,j*+1_); (3) if *c*_*i,j*_ ∈ *A* and *c*_*i,j*+1_ ∈ {−} then we need to keep the character *c*_*i,j*_ “saved” and continue going through the matrix until we reach a column *j* + *x* where 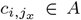 and *x* is some positive integer and *x >* 1, then we can draw an edge between *node*_*j*_ (*c*_*i,j*_) and *node*_*j*+*x*_(*c*_*i,j*+*x*_). An example of this final case in Figure 1 in the second sequence, the 5th column has a gap but the 4th has a *T*, we keep track of this until we reach the character *M* in the 7th column, where in that iteration we have constructed a node for the character *M* in column 7 and we can draw an edge between *node*_5_(*T*) and *node*_7_(*M*).

**Fig. 1.**
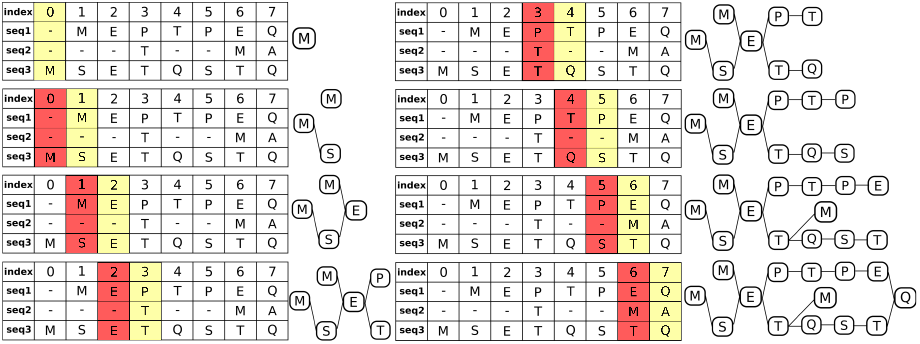
MSA to GFA: turning an MSA into a graph. The MSA in this example contains 3 sequences, - *MEPTPEQ*, - - - *T* - *MA*, and *MSETQSTQ*; and the step-by-step graph construction is shown on the panels from top to bottom. In every step, the yellow column is the current position and the red column is the previous one.

Because we iterate through the MSA from left to right and draw edges between consecutive nodes, the resulting graph is acyclic and directed.

#### 2.2.2 Compacting the graph

Linear stretches of nodes can arise while generating a graph from an MSA. Where the set of consecutive nodes 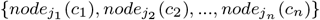 is a linear stretch of nodes if and only if each node in the set has an in-degree and out-degree of one, except the first node *node*_*j*1_ (*c*_1_) can have a higher in-degree and the last node *node*_*jn*_ (*c*_*n*_) can have a higher out-degree, then we can compact these nodes in the set into one node and concatenate their sequences. For example, in Figure 1 we see in the last step of constructing the graph, we have the stretch of nodes *P* → *T* → *P* → *E* that can be compacted into one node. `PanPA`’s final output is the compacted version of the graph in GFA format with each original sequence as a path entry in the output GFA.

##### Algorithm 1

Constructing a DAG from MSA

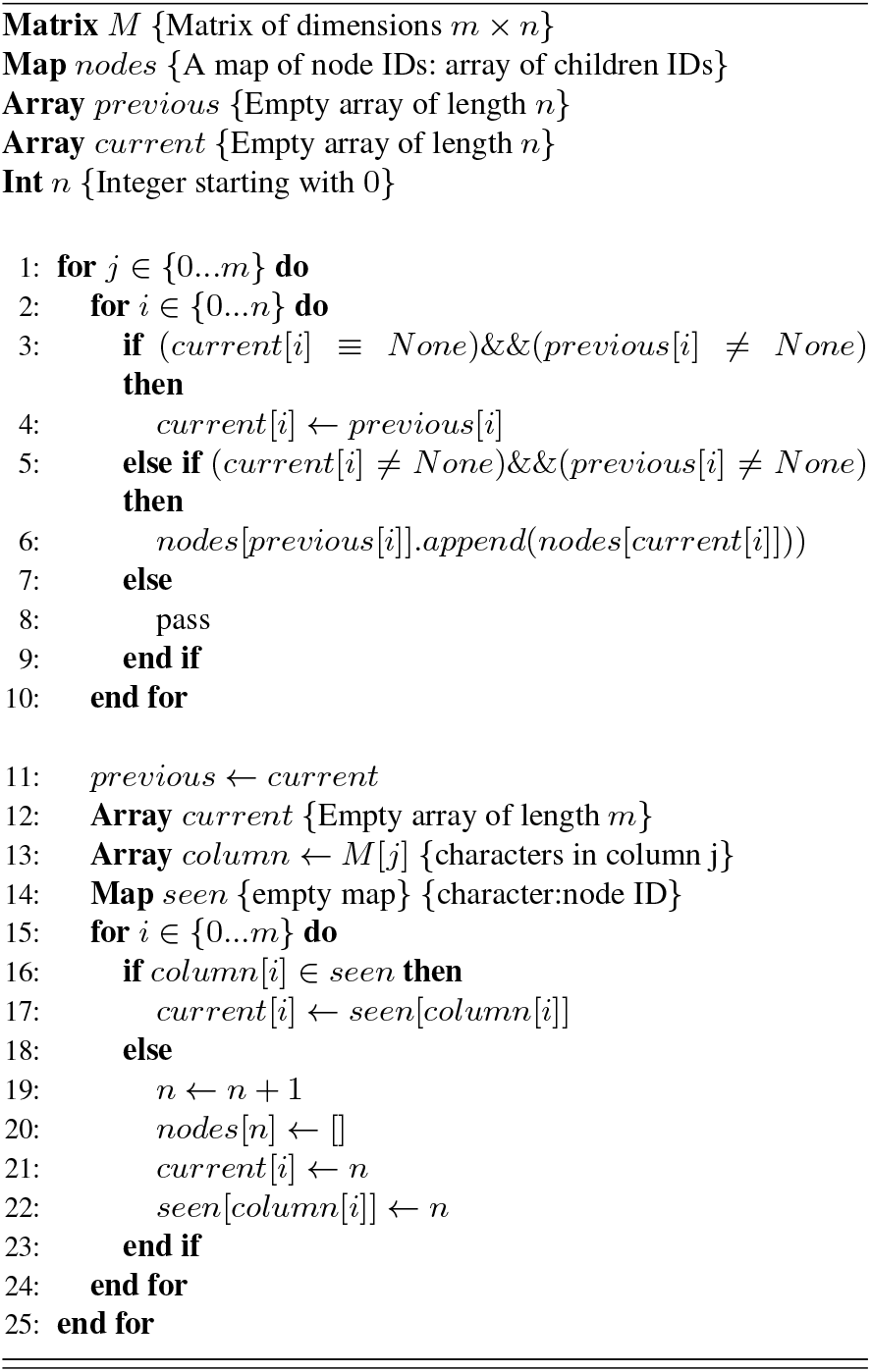

### 2.3 Aligning query sequences

`PanPA` uses a modified version of the known Smith-Waterman algorithm for local alignments (Smith and Waterman, 1981) known as partial-order alignment (Lee *et al*., 2002). The main idea of the modification is that instead of looking at the previous character in the alignment to fill the Dynamic Programming (DP) table, we need to consider all incoming edges of a node.

As each graph constructed from an MSA is a DAG, the graph can be topologically sorted, generating a list of ordered vertices. The concatenation of the sequences of the ordered vertices would be the target sequence to align against. As an example in Figure 2, the DAG and its concatenated sequence are shown as the columns in the dynamic programming matrix, and the first column contains the query sequence characters.

**Fig. 2.**
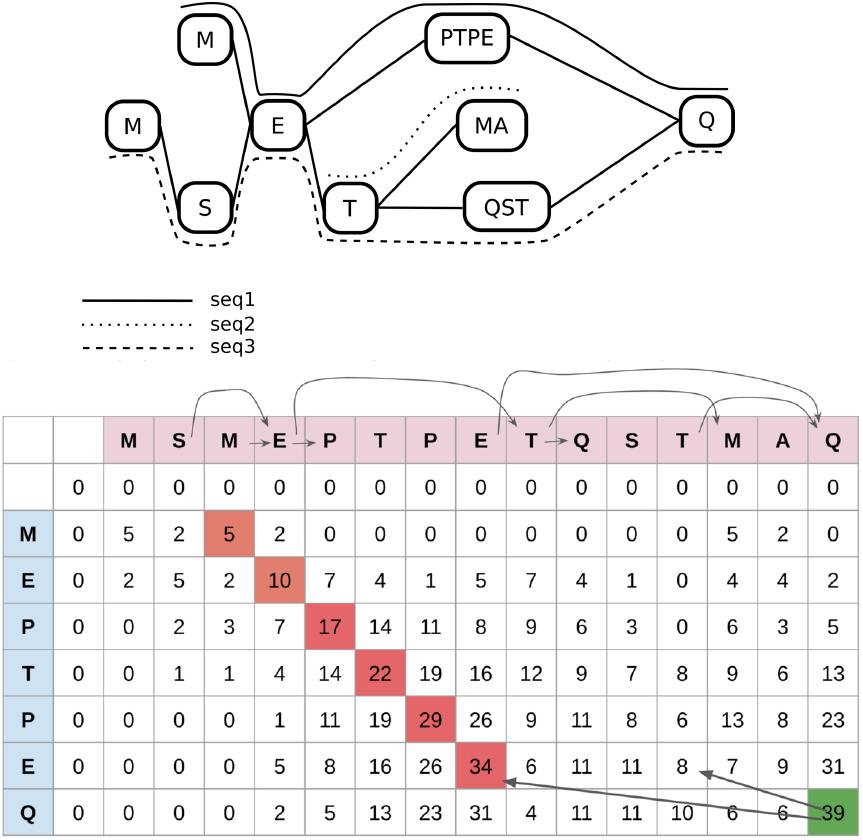
Alignment of a sequence to a protein graph. Top: example protein graph; bottom: the corresponding DP table. The ordered graph vertices are in the columns, the aligned sequence in the rows. Arrows between columns correspond to the graph edges. Arrows in the DP table correspond to potential previous cells in the DP process.

The dynamic programming matrix is defined as *H* = (*a*_*i,j*_) ∈ ℝ ^(*n*+1)*×*(*m*+1)^ where *m* is the size of the query sequence *M*, and *n* is the size of the concatenated sequences *N* from the ordered vertices. We add one extra row and column filled with 0 as the initializing row and column.

Similar to the Smith-Waterman algorithm, we need to fill the cells of the DP table using the information from the previous cells, where we usually look at the previous character. However, as some columns correspond to the first character of a node in the graph, we need to calculate the score of that cell based on all possible previous characters following all incoming edges to that node, as shown in Equation 1 for calculating the score of the cell *i, j*, we take the max of all scores calculated from all potential characters from the incoming edges *p*_*l*_ where *p*_*l*_ is the column index pointing to the previous character after following the incoming edge. To calculate a single score, Equation 2 is used, where we have 3 possible choices, a match/mismatch, an insertion, or a deletion, *δ* is the gap score, and *sub*(*c*_1_, *c*_2_) is a function that takes two characters and return the score based on a substitution matrix, e.g. Blosum62 (Henikoff and Henikoff, 1992).

Because our graphs are compacted, one node can have several characters. Therefore, if we are calculating the score for some *H*_*i,j*_ and the column *j* does not correspond to the first character in the node, we can simply then use Equation 2 and *p*_*l*_ is then simply *j* − 1.

For tracing back our alignment, we use the same approach of checking where the score of that cell came from to know which path our query sequence aligns to. For example, looking at Figure 2 at the last column corresponding to the character Q, if we start tracing back we see we have two incoming edges, one leading to the character E and the other to the character T, the score is then calculated for each previous character and the maximum score is taken, which lead us to E at *j* = 9, and we continue the traceback from there. We see for example that at *j* = 9 there are no incoming edges, so we only need to look at *j* = 8, and so on.

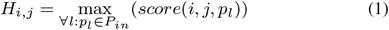

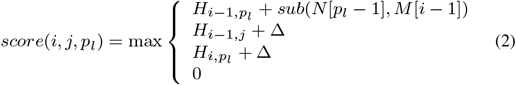

## 3 IMPLEMENTATION

`PanPA` was built using `Cython` without any extra dependencies, and `Cython` was used mainly to optimize the core alignment algorithm. Each step is implemented as a separate subcommand, facilitating it for the user to run each subcommand separately, which can help in case some parts needed to run more than once to find the optimal parameters for a certain dataset. The subcommands are *build_index, build_gfa*, and *align*.

Figure 3 shows an overview of `PanPA`’s workflow and the three key steps. It starts with MSA files, where each MSA represents one protein or a protein cluster. This input is accepted by both `build_index` and `build_gfa` modules. The subcommand `align` takes a FASTA file with query sequences, the graphs, and the index file produced from the `build index` step. `PanPA` then outputs the alignment in GFA format (Graph alignment format).

**Fig. 3.**
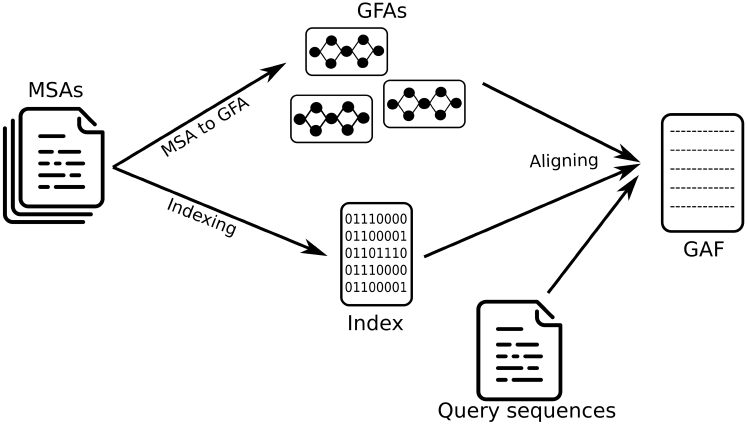
The general `PanPA` pipeline and its subcommands (in blue). Each subcommand can be also run separately or more than once with different parameters.

### 3.1 Indexing

In the indexing step, `PanPA` goes through each sequence in each MSA given and extracts the seeds from that sequence, be it *k*-mers or minimizers, depending on the user’s choice. Each seed is a key in a key-value map, and the value is a list of the MSA IDs where that seed was found. In our implementation, the value vector is ordered based on the number of times that seed showed up in that MSA normalized for the number of sequences in the MSA. Therefore, the user can then choose a cutoff limit on how many MSAs (equivalently, graphs) one seed can belong to, as some seeds can be promiscuous, especially when using a small value for *k*. Because the vector of hits is ordered, if the limit is an integer *n*, only the *n* top MSAs will be kept in the index.

For example, if we have three MSAs with IDs *m*_1_, *m*_2_, and *m*_3_ containing 10, 7, and 3 sequences respectively, and a seed *s*_1_ was present in *m*_1_ 2 times, in *m*_2_ 4 times, and *m*_3_ 3 times (with normalized counts being 0.2, 0.57, and 1 respectively), and the user cutoff is set to 2, then in our index, the seed *s*_1_ will point to the list [*m*_3_, *m*_2_].

In order to make extracting the minimizers from the consecutive windows faster, we used the Sliding Window Minimum algorithm from Carruthers-Smith (2011) which has a time complexity of 𝒪(*n*) where *n* is the size of the input sequence. This algorithm has also been described in more detail with pseudocode in Algorithm 1 from Jain *et al*. (2020).

### 3.2 Generating graphs

`PanPA` generates one corresponding DAG for each MSA in GFA format. Therefore, when a seed in the index points to one MSA, we can align the query sequence to the graph that corresponds to that MSA. Moreover, because the original sequences in the MSA are encoded as paths with the path line in GFA, we cannot compact two adjacent nodes connected by one edge if not the same set of sequences pass through both these nodes. For example, imagine we have an MSA with the following 3 sequences *MT QT*, - - *QT*, and *MT* - -. This will generate a graph with a linear stretch of 4 vertices (*M, T, Q*, and *T*) with one edge between every two consecutive vertices. However, if we compact all four vertices into one, we cannot write a path for sequences 2 and 3 in the GFA file, because now they are contained inside this compacted node. Therefore, we can only compact nodes *M* and *T* together and nodes *Q* and *T* together, this way sequence 3 is contained in the first node and sequence 2 in the second. Supplementary Figure 3 visualizes this example and shows the corresponding GFA files for the graph before and after compacting.

### 3.3 Aligning

Given a query sequence, we count all the seed hits from the query to the MSAs using the index and generate a list of MSAs (equivalently, graphs) to align against, the list is sorted based on the number of hits: for example, if the query sequence had 5 seeds, where 4 of them pointed to *m*_1_, and one pointed to *m*_3_, our list of matches will be [*m*_1_, *m*_3_], the user can also specify to how many potential MSAs/graphs can one query be aligned against, or choose to align to all matches, if, for example, the limit of the number of seeds was set to 1, our query sequence will only be aligned to *m*_1_. Moreover, the user can specify a minimum acceptable alignment identity score, and only the alignments with scores equal or larger to this minimum threshold are returned. `PanPA` also uses a linear gap penalty and the user can choose one of many substitution matrices available.

## 4 RESULTS

### 4.1 Validating PanPA on a panproteome of *E. coli*

We first wanted to validate that `PanPA` is able to find correct alignments. Therefore, we applied it to an *E. coli* panproteome we built. To that end, we first downloaded 1,351 *E. coli* assemblies that were marked as “Complete Genome” from RefSeq (O’Leary *et al*., 2016). We extracted every amino acid sequence corresponding to a coding region from the annotations provided on RefSeq and proceeded to cluster these sequences using `mmseq2` (Hauser *et al*., 2016) with default parameters, this resulted in 44,204 protein clusters. The distribution of the number of strains per cluster (Supplementary Figure 1) has the characteristic U-like shape which evidences the presence of the core genes that are present in nearly all assemblies (right part of the plot) and accessory genes that are mostly unique to one assembly or present in only a few (left side of the plot). Now that we had similar proteins clustered together, `mafft` (Katoh and Standley, 2013) was used on each cluster to produce a corresponding MSA.

We then proceeded with `PanPA` to produce a DAG in GFA format from each MSA. We randomly selected 32,289 protein sequences from our MSAs collection. The random selection was done by, first, randomly selecting 10% of all the MSAs representing the protein clusters, then for each MSA chosen, we randomly selected 5% of sequences in that MSA. Therefore, we have a ground truth as to where each sequence comes from and to which graph it should align against; and we expected that `PanPA` should align each of these sequences to the correct corresponding graph. We constructed a pipeline using `Snakemake` (Mölder *et al*., 2021) to run the indexing and alignments steps with a combination of several parameters to demonstrate the effect of different parameters on the correctness of the alignments.

We define a “wrong alignment” here when the highest-scoring alignment produced by `PanPA` corresponds to an alignment against a different graph/MSA than that where the sequence originated from. One can observe in Figure 4 that for *k* = 3 we get a relatively high number of wrongly aligned sequences, unless the index stores all the seed hits (the value 0 in the figure, with the red marks), whereas higher *k* values produce very few wrong alignments regardless which cutoff was used for the index. Moreover, using indexes with small *k* and *w* values also results in higher alignment time as more seeds need to be extracted, the seeds have more matches, and more look-ups need to be done to find the top potential graphs to align back to. For example, in this experiment, taking *k* = 3 will require a maximum of 659 seconds in system time, whereas it takes less than 100 seconds using *k* = 9; Supplementary Figure 2 shows the system time for all combinations of parameters. From these results, we can recommend that a *k* value bigger than 3 should be used for alignments against closely-related species, and the user can use a cutoff on the index without losing too many alignments. For a full sensitivity, we recommend that the user uses a small *k* value but should not limit the index and keep all seed hits. However, this will result in a longer alignment time.

**Fig. 4.**
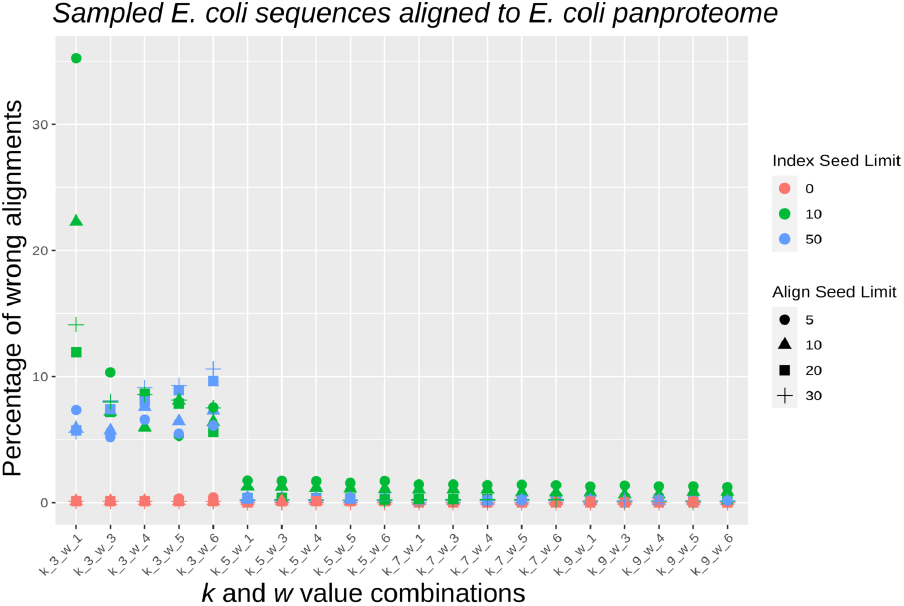
This figure demonstrates the effect of the different parameters with respect to the wrongly aligned sequences, where a “wrong alignment” is a sequence being aligned to a different graph than the one it originated from. Each point is also colored based on the seed index limit (the limit of how many hits can each seed point to), and the shapes correspond to the alignment seed limit (the limit of how many graphs can one sequence align against). We see that for small *k* values, we get a high number of wrong alignments unless when not limiting the index size. The align seed limit has a relatively small effect on the wrong alignments

### 4.2 Aligning unseen sequences in *E. coli*

Using the same panproteome constructed in the previous experiment, we further downloaded 80 *E. coli* assemblies from RefSeq that were not used in building the panproteome and extracted the protein sequences from the corresponding annotation files. After removing redundant sequences, we were left with 92,196 sequences. We used the same `Snakemake` pipeline as in the previous experiment to align these sequences against the panproteome with the different parameter combinations. In Figure 5 we show the effect of the different parameters on the number of unaligned sequences, where alignments had to pass the 90% identity threshold. We observe again that for small values of *k*, the majority of sequences (between 50% for *k* = 3 and *w* = 6 to 99% for *k* = 3 and *w* = 1) did not produce an alignment. These results emphasize the conclusion from the previous experiment, that choosing a very small size for the seeds (e.g. *k* = 3) will result in a high number of false positive index hits that result in alignment with low identity and get filtered out, especially when the index is limited. Moreover, when the index is unlimited, then `PanPA` is able to find the correct graphs. However, an unlimited index will result in much higher alignment time as there is a need to align to more sequences, for example, for *k* = 3, *w* = 1, and unlimited index, it takes `PanPA` over 3285,4 seconds of system time to finish alignments compared to less than 402 seconds with *k* = 9 and *w* = 1. Supplementary Figure 3 is similar to Figure 5 but with system time as the y-axis.

**Fig. 5.**
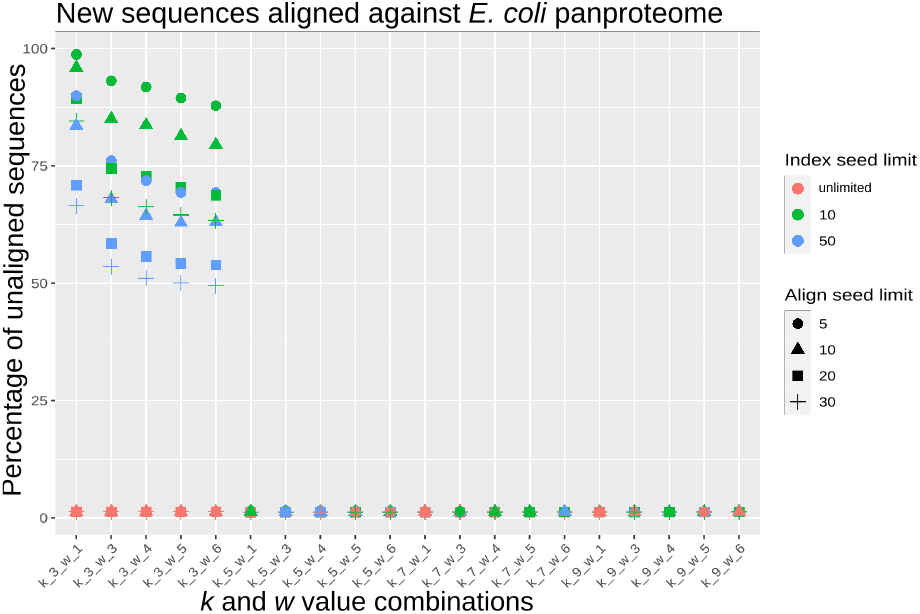
This figure shows the effect of the different parameters on the number of unaligned sequences when aligning the new 92,196 *E. coli* sequences. We observe that for small *k* values, the majority of sequence were not aligned, unless we do not set a limit for the index (the red marks); then over 99% of sequences produce an alignment.

### 4.3 Comparison of PanPA with BWA and GraphAligner using *S. enterica* sequences

One of the major advantages of moving to the amino-acid space is the ability to have better alignments between more distant organisms. To test this, we downloaded 1,077 *Salmonella enterica* annotated assemblies from RefSeq, extracted all coding regions and aligned them to the *E. coli* assemblies and graphs that we have already. Both *E. coli* and *S. enterica* belong to the same family *Enterobacteriaceae*, but are from different genera, and hence are expected to be far apart from each other evolutionary to make a good test case for our tool `PanPA`. In order to compare DNA and protein alignments, we extracted all DNA sequences of coding regions and their corresponding amino acid sequences from the *S. enterica* annotations, we obtained 4,839,981 sequences, which we used to align to *E. coli*.

We compared three kinds of alignments here: (1) DNA sequence alignments against the *E. coli* linear reference genome (strain K-12 substrain MG1655) using `BWA` (Li and Durbin, 2010); (2) DNA sequence alignments using `GraphAligner` (Rautiainen and Marschall, 2020) against the *E. coli* pangenome graph from all 1,351 assemblies that we constructed with `minigraph` (Li *et al*., 2020); and (3) amino acid sequence alignments using `PanPA` against the *E. coli* panproteome constructed in the first experiments. Both `BWA` and `GraphAligner` were run with default parameters, and `PanPA` was given an index with *k* = 5, *w* = 5, an index limit of 10, and only aligning each sequence to the top 10 graph hits. The alignments were then filtered based on alignment length and alignment identity, and only alignments with a length of over 50% of the original sequence length and alignment identity of at least 50% were kept.

Out of the 4,839,981 sequences, 1,638,936 were successfully aligned by all three aligners, while 1,694,181 could only be aligned by the graph-based methods GraphAligner and PanPA. Strikingly, `PanPA` could align 744,033 unique sequences that were not aligned by any of the other two aligners (Figure 6). `PanPA` alignments have higher identity scores, which is to be expected as in the amino acid space the sequence identity is higher for the same two sequences as in the DNA space (Figure 7). This confirms the advantages of aligning using the amino acid alphabet, which PanPA now enables to leverage for sequence-to-graph alignments. Numbers corresponding to all combinations in Figure 6 can be found in Supplementary Table 1.

**Fig. 6.**
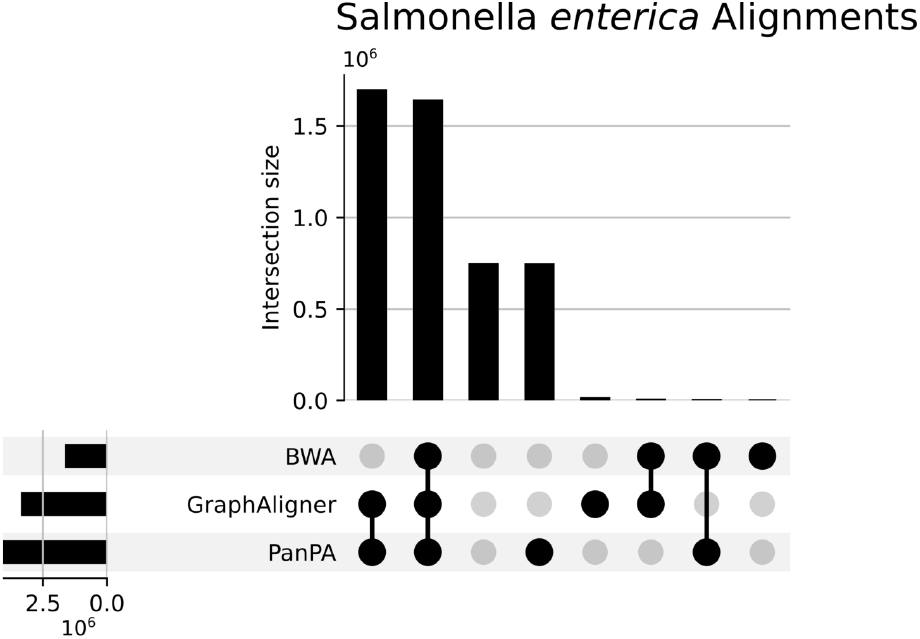
Upset plot of the unique alignments of 4,839,981 sequences from the coding regions of 1,074 *S. enterica* assemblies from RefSeq. Alignments with `BWA` and `GraphAligner` (DNA), and `PanPA` (amino acids) against their corresponding *E. coli* counterparts were constructed using the parameters in Supplementary materials section 2.

**Fig. 7.**
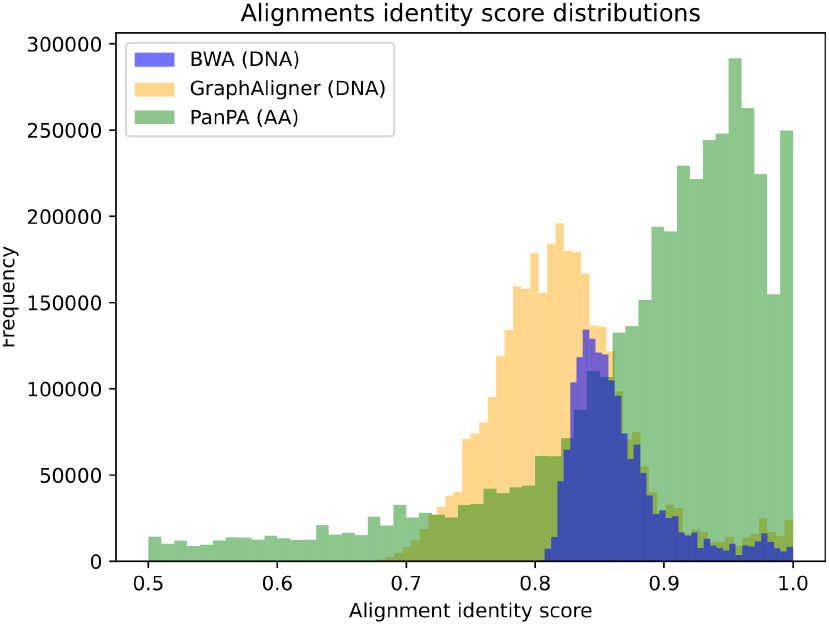
Distribution of identity scores between `BWA`, `GraphAligner`, and `PanPA` from aligning the *S. enterica* sequences. We see that the pique for `PanPA` is shifted to the right, which is expected, as amino acids have higher identity compared to nucleotide sequences

### 4.4 Aligning *S. enterica* Illumina short reads against *E. coli*

`PanPA` is also able to align DNA sequences to protein graphs by translating each DNA sequence into 6 different reading frames (3 forward and 3 reverse-complement). This feature can be very helpful for aligning sequencing reads from organisms that do not have a reference or a close-enough reference to align against.

We downloaded one *S. enterica* Illumina whole genome sequencing (WGS) short reads sample (SRR22756191) from NCBI SRA database (Leinonen *et al*., 2011) containing 1,110,471 sequences, which is part of PulseNet USA surveillance for foodborne disease. We aligned the sequences using `BWA` against the linear reference of *E. coli* that we used in the previous experiment and against the *E. coli* panproteome using `PanPA`, we used the index with *k* = 5, *w* = 3, and no cutoff on the index. In the alignment step, we allowed each sequence to align to up to 20 graphs. We filtered the output alignment based on 50% alignment identity score and 50% alignment length. `BWA mem` was used with default parameters. We chose a relatively small *k* value because we wanted higher sensitivity, and because every read is translated into 6 different reading frames, there is a higher chance of false positive hits when finding hits for the sequences in the index. However, increasing the number of allowed graphs will only affect the number of alignments done and the overall run time, in the end, the alignments that do not pass the identity threshold will not be returned. Similar to the experiments above, we advise choosing a smaller seed size with an unlimited index when the user wants higher sensitivity, like aligning against a more distant organism.

The results in Table 1 show that using a distant linear reference has a major disadvantage as around 65% of the reads did not align with identity over 50% and after filtering based on both alignment length and identity being over 50%, only 4.4% were reported by `BWA`. On the other hand, `PanPA` was able to produce alignments for 97% of the sequences with identity over 50% and 69% of sequences had an alignment after further filtering based on the alignment length. There were 334,414 sequences that were not aligned from either aligners after filtering.

**Table 1.** Number of *S enterica* DNA short reads aligned against *E. coli* ‘s linear reference with `BWA` and against its panproteome using `PanPA`.

| | Identity ≥ 50% | Identity and length ≥ 50% |
| --- | --- | --- |
| BWA | 391,041 (35.2%) | 48,937 (4.4%) |
| PanPA | 1,080,315 (97.3%) | 768,264 (69.2%) |

### 4.5 Using PanPA to display phenotypic traits: a case of antimicrobial resistance in *E. coli*

Certain mutations are associated with bacteria being resistant or susceptible to antibiotics, and this has been a main focus of many researchers as resistance against antibiotics presents a major threat to public health. We explored the applicability of our tool `PanPA` to identify such mutations. To this end, we used the Pathosystems Resource Integration Center (PATRIC) (Davis *et al*., 2020) database which contains assemblies and annotations for many antibiotic-resistant and susceptible bacteria. We downloaded the ciprofloxacin-resistant and -susceptible strains from *E. coli*, which comprised a set of 1,295 susceptible and 507 resistant genome assemblies. From these assemblies and their annotations we extracted two genes, *parC* and *gyrA* which encode for quinolone, and particularly ciprofloxacin, targets and can carry resistance-associated mutations in *E. coli* (Bagel *et al*., 1999). For each of these two proteins, we randomly split the sequences into two sets, one containing 10% of the sequences and the other 90% (46+461 for resistant bacteria, and 117+1295 for susceptible) we then mixed the 90% part of both susceptible and resistant together for each protein, generated an MSA for each using `mafft` and then a graph for each protein respectively using `PanPA`(). We visualized the resulting graphs in Figure 8. First, nodes in the graph were colored depending on the percentage of resistant and susceptible sequences passed through these nodes, which made it clearly visible that certain mutations differentiate between these two states, and thus are likely associated with resistance. Second, we aligned the 10% sequence set back to the graphs using `PanPA` and draw different lines in Figure 8 showing which paths the alignments followed in the graph, we also notice that the vast majority of the sequences extracted from resistant strains aligned to the nodes that seem to be associated with resistance, and susceptible sequences aligned to mostly nodes associated with susceptible strains.

**Fig. 8.**
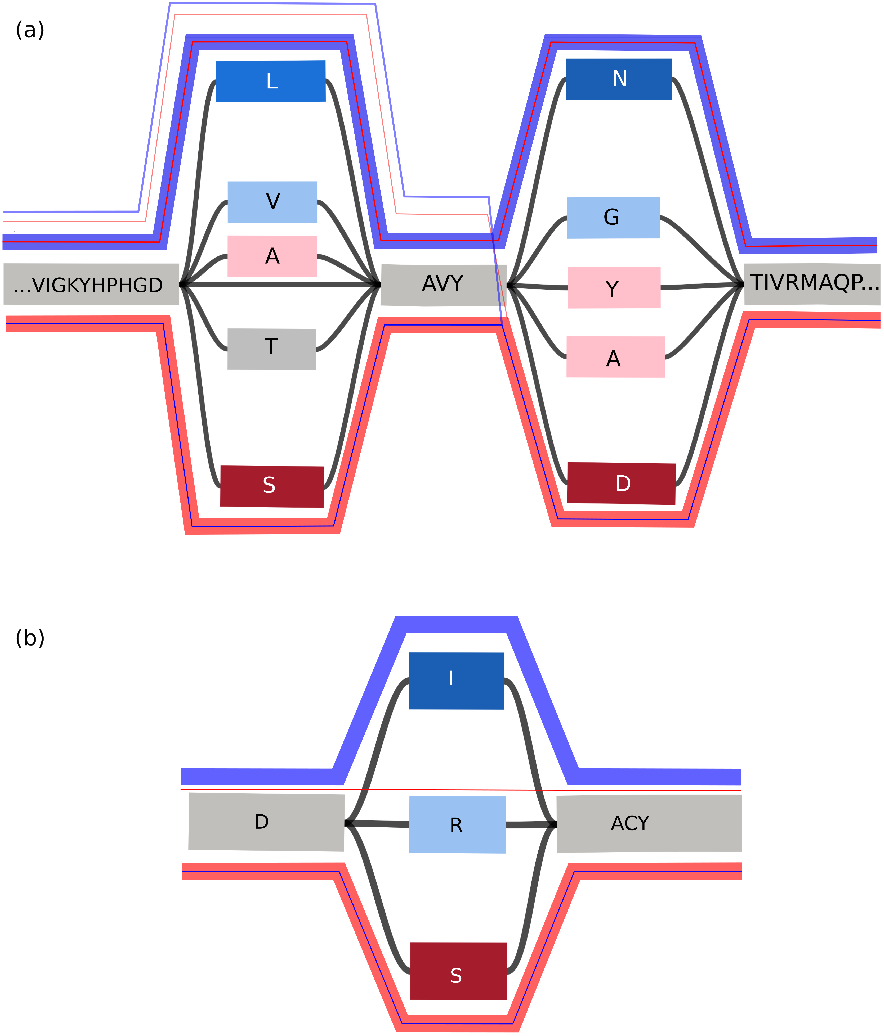
Visualization of parts of the protein graphs for (a) GyrA and (b) ParC using `Bandage` (Wick *et al*., 2015). Nodes are colored depending on the number of resistant/susceptible strains that pass through them, the blue color represents resistance, and red represents susceptibility, the intensity of the node’s color correspond to the number of susceptible or resistant sequences passing through these nodes. The lines on top of the graph represent the paths that the 10% sequence set aligned (45 resistant sequences and 117 susceptible sequences) took, the color represents the type, and the thickness represents the number of sequences taking that path. We see a thick blue line of resistant sequences took the blue path passing through the blue nodes, and vice versa, a thick red line for susceptible sequences took the red path passing through the red nodes.

In this case, we observe well-known resistance-associated mutations Ser83 and Asp87 in GyrA (Webber *et al*., 2017; Rakici *et al*., 2021; Yu *et al*., 2020) and Ser80 in ParC Nawaz *et al*. (2015) that preclude drug binding on the protein-DNA interface. Besides the canonical resistance-associated variants (S83L, D87N in GyrA and S80I in ParC), we observe other potential variants that are present predominantly in resistant strains: alanine, leucine and valine at position 83 and alanine, tyrosine, and asparagine at position 87 of GyrA, as well as arginine at position 80 of ParC.

## 5 CONCLUSION

In this paper, we present `PanPA`, a software tool to build and index panproteome graphs, and align sequences to them. In our method, instead of building one big graph that represents all samples of a population, we build many local graphs, where each local graph represents one protein or a group of related proteins.

We demonstrate that `PanPA` produces correct alignments when aligning a sample of *E. coli* protein sequences back to an *E. coli* panproteome produced from assemblies from a public database. We also show that moving into amino acid space can increase both the number of aligned sequences and the alignment identity score when comparing phylogenetically distant organisms as exemplified by aligning *Salmonella enterica* proteins against the panproteome constructed from *E. coli* assemblies. Also showing that `PanPA` can capture a much higher number of DNA sequence alignments that would have been otherwise lost when using a distant reference. We argue that aligning over longer phylogenetic distances is important, especially when trying to study organisms that are not well-researched, do not have a standard reference, and where a particular clade is only scarcely sequenced. In these cases, one can use a distant organism panproteome to produce better alignment and comparison, maybe one step towards annotation using a remote reference. Additionally, we demonstrate the utility of `PanPA` for the discovery of genetic mechanisms of phenotypic traits, such as antimicrobial drug resistance.

## 6 AUTHOR CONTRIBUTIONS STATEMENT

F.D, T.M, and O.K conceived the study. F.D wrote `PanPA`, ran experiments, and wrote a draft of the manuscript. S.S contributed to part of the code. T.M and O.K supervised the work and edited the manuscript. All authors read and approved the final version of the manuscript.

## ACKNOWLEDGEMENTS

The authors would like to thank Konstantinn Bonnet and Sebastian Keller on code review and discussions, and Amay Agrawal for helping in obtaining the PATRIC resistant bacteria data.

