## supplementary_material for "PanPA: generation and alignment of panproteome graphs"

### Supplementary Material
### PanPA: generation and alignment of pan-proteome graphs

January 19, 2023

| Intersection | Number of alignments ≥ 50% identity | Number of alignments ≥ 70% identity |
|---|---|---|
| Not Aligned | 744,964 | 1,012,744 |
| BWA | 1 | 1 |
| BWA - GraphAligner | 4,084 | 4,090 |
| BWA - PanPA | 1,294 | 1,294 |
| GraphAligner | 12,488 | 52,357 |
| Graphaligner - PanPA | 1,694,181 | 1,643,479 |
| PanPA | 744,033 | 487,086 |
| BWA - GraphAligner - PanPA | 1,638,936 | 1,638,930 |

Supplementary Table. 1: Intersection of unique alignments of 4,839,981 sequences representing the annotations from Salmonella *enterica* assemblies from RefSeq, against E. *coli* linear reference, pangenome, and panproteome using BWA, GraphAligner, and PanPA respectively.

| Aligner | BWA | GraphAligner | PanPA |
|---|---|---|---|
| **Num. alignments** | 2,699,361 | 26,009,077 | 8,684,414 |
| **Num. filtered alignment 50% length** | 1,645,224 | 4,399,906 | 7,897,707 |
| **Num. filtered alignments 50% id** | 1,645,224 | 4,399,906 | 7,897,707 |
| **Num. filtered alignments 70% id** | 1,645,222 | 4,384,913 | 5,273,200 |

Supplementary Table. 2: Number of alignments from the 4,839,981 sequences from Salmonella *enterica* annotations using BWA, GraphAligner and PanPA. We see that GraphAligner produced the most alignments. However, after filtering for an alignment length of at least 50% of the original sequence size, the number of alignments drops drastically. For PanPA, most of the alignments were long enough and only a small number got filtered.

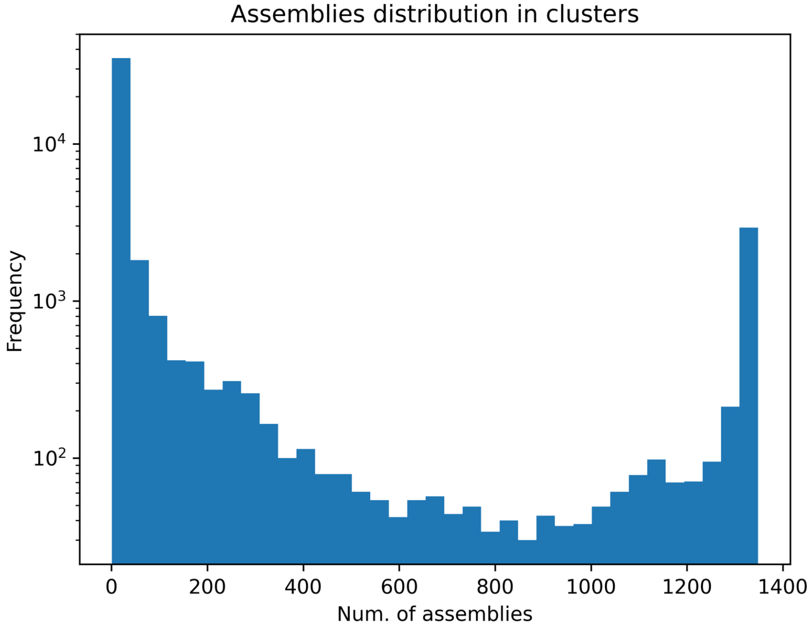


Supplementary Figure. 1: Histogram of number of different strains in each cluster, the typical U-shape cluster where the left peak represent the unique clusters, and the right peak represent the core genes cluster

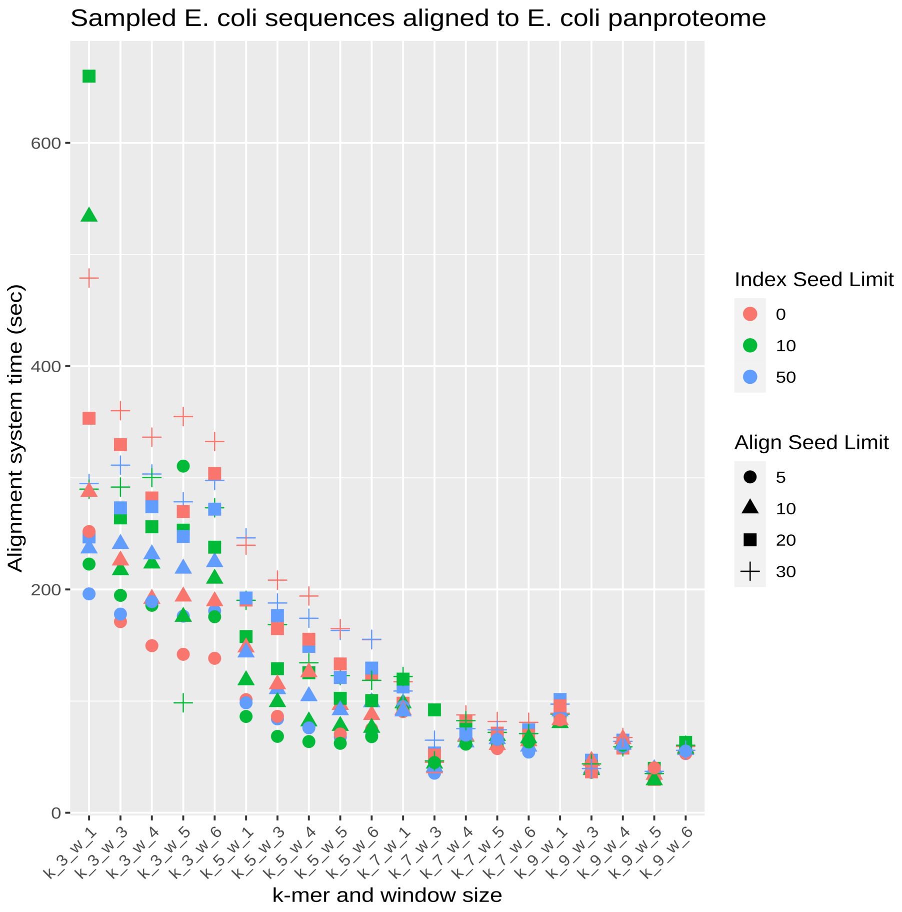


Supplementary Figure. 2: $k$ and $w$ sizes against alignment time. We can see the indexes with smaller values for $k$ and $w$ will have higher alignment time.

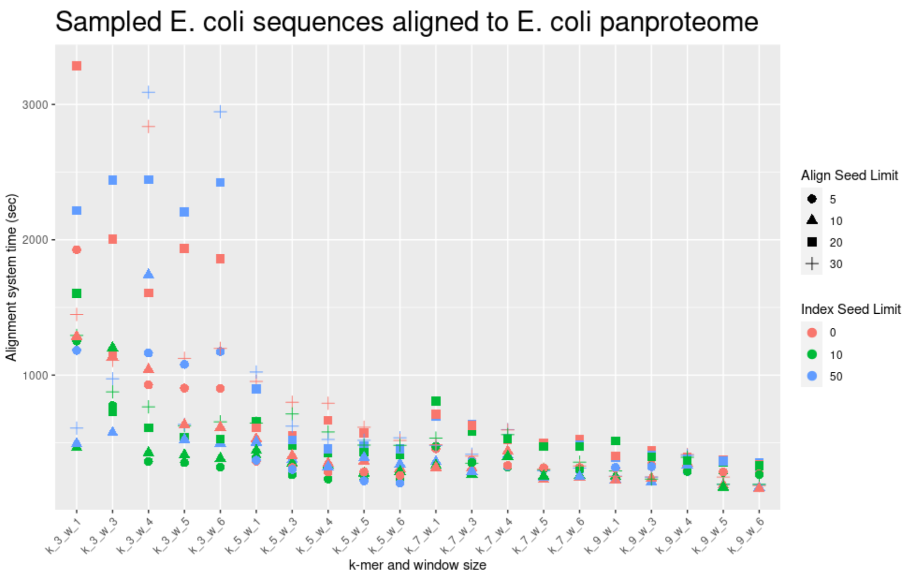


Supplementary Figure. 3: $k$ and $w$ sizes against alignment time for the unseen *E. coli* sequences aligned against the panproteome. We see that for small values of $k$ the time is much higher, especially when the index is unlimited.

### 1 Sequences Random Selection

The 32,289 sequences from the *E. coli* panprotome that were chosen for testing were chosen at random using a script that can be found on `PanPA`'s repository, it takes two arguments as input, both integers from 0 to 100 representing the percentage of different protein clusters to choose at random, and how many sequences to choose from each MSA. We gave the script the inputs 10 and 5, which then chooses at random 10% of the protein clusters, and then from each cluster chooses 5% of the sequences at random. Because we know from which cluster each sequence belongs to, we can calculate the number of matches after doing the alignments.

### 2 *S. enterica* Alignments Parameters

For aligning the DNA sequences from *S. enterica* against the *E. coli* reference genome, we use `BWA` with the following parameters:

```
bwa mem e_coli_reference_GCF_000005845.2_ASM584v2.
    fasta salmonella_refseqdna.fasta -t 60 >
    salmonella_refseqdna_ecoli_ref_genome_bwa.sam
```

We used `GraphAligner` for aligning *S. enterica* DNA sequences against *E. coli* pangenome that was built using `minigraph` with the following parameters:

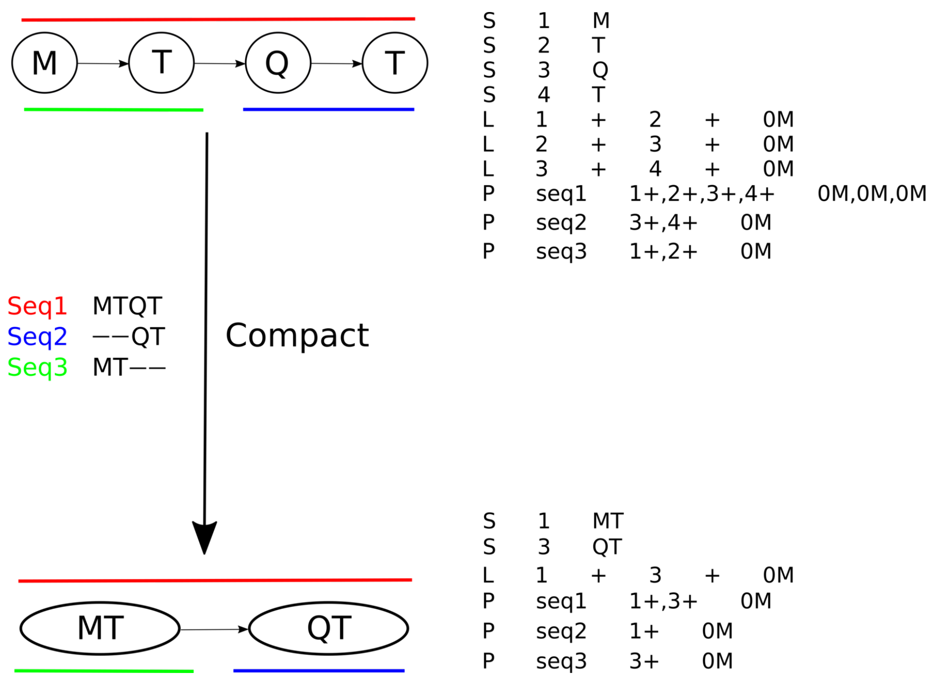


Supplementary Figure. 4: An example of constructing a graph from an MSA with 3 sequences $MTQT$, $QT$, and $MT$. The top left graph is the uncompacted graph, where each column in the MSA produced one node, the red line is the path for Sequence 1, the blue for sequence 2, and the green for sequence 3, to the right of the graph is the GFA format that would produce that graph with these paths for each sequence. After compacting, instead of ending up with only one node as we have have a path graph, we end up with two nodes, because in the GFA format, a path is represented with nodes, and if we compact all the nodes, we will not be able to represent Sequence 2 and 3. Therefore, nodes 1 and 2 get merged, and 2 and 3 are merged

```
GraphAligner -f salmonella_refseq_dna.fasta -g
    e_coli_pangenome.gfa -a
    salmonella_refseqdna_ecoli_pangenome.gaf -x vg --
    threads 60 2> graph_align.log

# for building the pangenome, these commands were used
# this command is the initial one to build a graph
minigraph -xggs -t20 e_coli_reference_GCF_000005845.2
    _ASM584v2.fasta e_coli_reference_GCF_000005845.2
    _ASM584v2.fasta > e_coli_pangenome.gfa

# updating the graph by adding one assembly every step
# assemblies_locations.txt is a list of each E. coli
    assembly to update the graph
while read r;do minigraph -xggs -t20 e_coli_pangenome.
    gfa $r > tmp && mv tmp e_coli_pangenome.gfa;done <
    assemblies_locations.txt
```

PanPA was used with the following parameters:

```
PanPA --log_file salmonella_aa.log align -d e_coli_gfa
    / --index index_k_5_w_5_seed_lim_10.pickle -r
    salmonella_aa.fasta.gz -o
    salmonella_aa_ecoli_panproteome.gaf --min_id_score
    0.5 --cores 50 2> panpa_time.log
```

#### 2.1 Aligning short reads parameters

For aligning the short reads sample of *S. enterica* from SRA database with accession number SRR22756191. The following command was used for BWA:

```
bwa mem -t 5 reference_ecoli_GCF_000005845/
    SRR22756191.fasta > alignments_SRR22756191.sam
```

For PanPA, the following command was used:

```
PanPA --log_file short_reads_log.log align -d
    e_coli_graphs/ --index index_k_5_w_3_seed_lim_0.
    pickle -r SRR22756191.fasta --dna -c 10 -o
    SRR22756191_ecoli.gaf --min_id_score 0.5 --
    seed_limit 20
```
